# Ecological dynamics and stability in the Taï and Comoé national parks in Côte d’Ivoire

**DOI:** 10.64898/2026.08.12.744377

**Authors:** Jean-Louis Kouakou, Atta Assemien Cyrille-Joseph, Yao Kouassi Alphonse, Amara Ouattara, Abdoulaye Diarrassouba, Séry Gonedelé-Bi

## Abstract

The accelerated loss of biodiversity in sub-Saharan Africa threatens the functioning of tropical ecosystems. In Côte d’Ivoire, the Comoé National Park (PNCOMOE), a Sudano-Guinean savannah, and the Taï National Park (PNTAI), a dense rainforest, both UNESCO World Heritage Sites, are home to fauna assemblages of global importance, whose long-term resilience remains insufficiently quantified. This study assesses and compares, over a decade (2014-2025), the functional stability of vertebrate communities in these two contrasting ecosystems, using nine metrics covering resistance, invariance, persistence, interspecific synchrony, Tilman’s stability, Jacobian resilience and a Composite Stability Index (CSI). Abundance data for 107 vertebrate species were collected via foot transects at PNTAI and aerial surveys at PNCOMOE. The stability metrics were calculated using the R package ‘estar’, integrated with an alpha diversity analysis (Shannon H′, species richness S, Pielou’s evenness J′) and a Jacobian spectral analysis within a multidimensional ecological assessment. PNTAI (0.708) exhibits significantly higher alpha diversity (H′ = 2.82; S = 55.7 taxa) and community resilience 4.6 times higher than in the PNCOMOE (0.153). Its interspecific asynchrony index (0.504) reveals a strong portfolio effect, absent in the PNCOMOE (0.232). In contrast, PNCOMOE exhibits higher temporal invariance (0.382 versus 0.116) and Tilman stability (0.276 versus 0.152), reflecting more predictable dynamics. The overall ICS favours the PNTAI (0.484) and (0.370). The Jacobian analysis detects local instability in both parks (Re(λmax) = 5.58 at the PNTAI; 3.73 at the PNCOMOE). The two parks exhibit distinct yet complementary stability architectures: PNTAI relies on dynamic stability based on resilience and interspecific compensation, whilst PNCOMOE demonstrates conservative stability through temporal regularity. The absence of calculable resilience at PNCOMOE suggests a potential crossing of a functional degradation threshold, arguing for urgent restoration interventions and differentiated conservation strategies, tailored to the resilience mechanisms specific to each ecosystem.

## Introduction

Protected areas are essential pillars of global biodiversity conservation, particularly in tropical regions where species richness, endemism and the functional complexity of ecosystems reach exceptional levels [1–3]. In West Africa, national parks play a major role in preserving natural habitats, protecting threatened species and maintaining fundamental ecological processes [4,5].

In Côte d’Ivoire, Taï National Park and Comoé National Park represent two of the most important protected areas in the country and the West African sub-region. Comoé National Park, a UNESCO World Heritage Site, is one of the largest protected areas in West Africa, covering an area of approximately 1,148,756 hectares [6–8]. It is characterised by an ecological transition zone between Guinean forest formations and Sudanese savannahs, thus offering a mosaic of habitats particularly conducive to high biological diversity [8].

Conversely, Taï National Park, also a World Heritage Site, represents one of the last significant tracts of primary tropical rainforest in West Africa [9,10]. It is home to remarkable biodiversity, including several iconic and threatened species such as *Pan troglodytes verus, Loxodonta cyclotis* and numerous species of duikers and forest primates [9,10].

Despite their ecological and heritage importance, these two ecosystems have been under significant pressure for several decades, notably from poaching, habitat fragmentation, climate change, peripheral agricultural activities and anthropogenic disturbances linked to local socio-economic dynamics [7].

In this context, assessing the temporal dynamics of biological communities and their ecological stability has become a major scientific challenge for the sustainable management of these protected areas. Ecological stability, a central concept in community ecology, refers to a system’s capacity to maintain its structure and functions in the face of disturbances, as well as its ability to recover after stress [11,12]. It can be understood through several complementary dimensions, notably resistance, resilience, invariance, persistence and dynamic stability [13,14].

A comparative study of the dynamics of abundance, alpha diversity, species richness, community evenness and functional stability metrics between Taï and Comoé National Parks thus provides a better understanding of the ecological mechanisms underpinning the robustness or vulnerability of these ecosystems.

The aim of this study is therefore to analyze the spatio-temporal dynamics of wildlife communities and to assess the ecological stability of the two main Ivorian national parks, to identify differences in functioning between savannah and tropical rainforest systems, and to provide scientific insights useful for their conservation and adaptive management.

## Materials and methods

### Study area

This study was conducted in two major protected areas in Côte d’Ivoire: the Comoé National Park (PNCOMOE) and the Taï National Park (PNTAIAI). These two sites were selected on account of their ecological importance and their representativeness of the two major West African tropical biomes (Fig 1). PNCOMOE, located in north-eastern Côte d’Ivoire, forms part of the Sudano-Guinean savannahs and is one of the largest protected complexes in West Africa, characterized by a mosaic of wooded savannahs, shrub savannahs, gallery forests and wetlands (Fig 1). The PNTAI, situated in the south-west of the country, in contrast, represents a massif of dense tropical rainforest that is among the best preserved in the sub-region, renowned for its exceptional biological richness [Fig 1]. The choice of these two contrasting ecosystems allows for a robust comparison of stability dynamics between a savannah system and a tropical forest system, in line with the comparative approaches recommended in community ecology [12,13].

**Fig 1.**
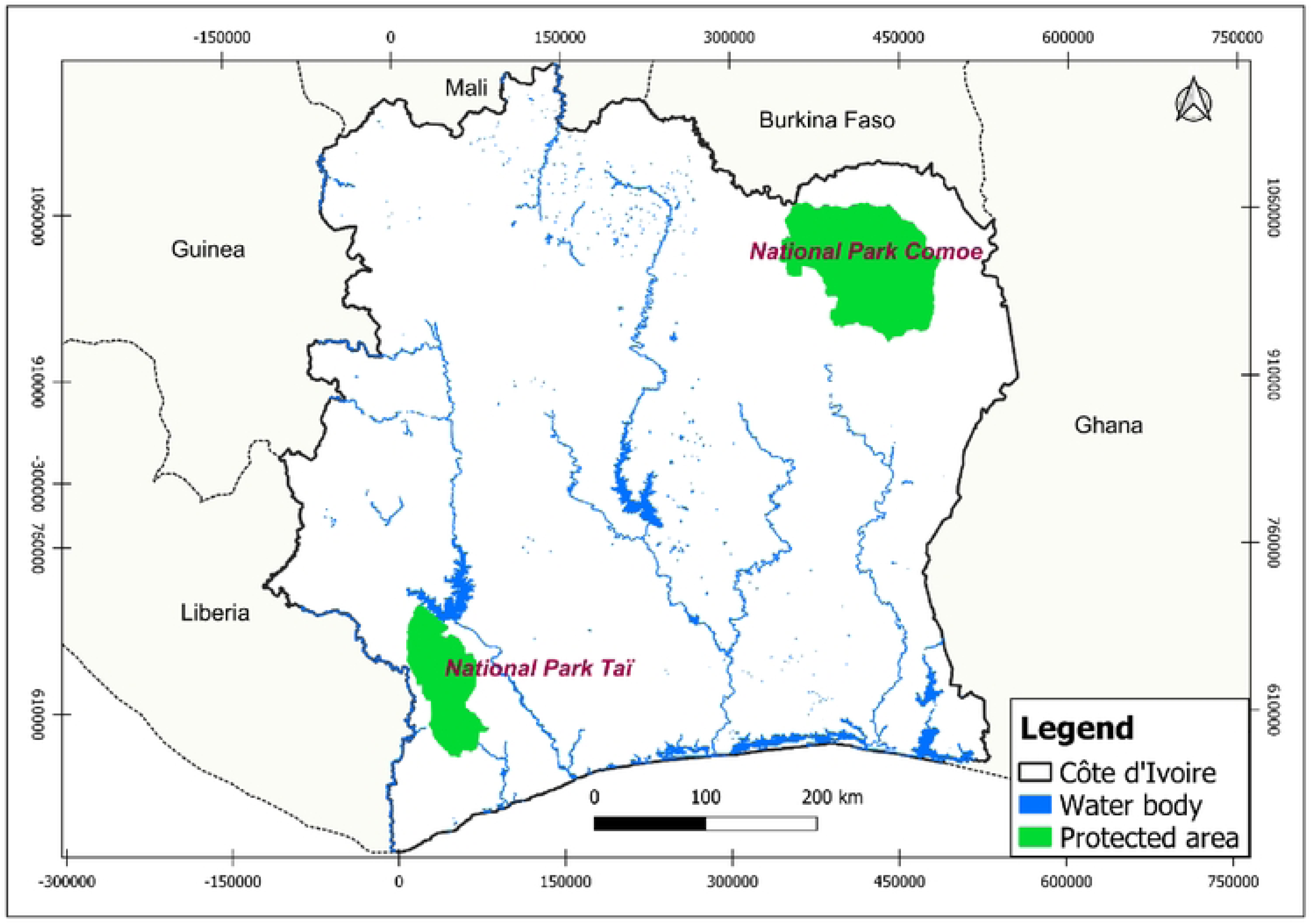
Study site.

### Data collection and organization

The data used were derived from wildlife surveys conducted between 2014 and 2025 by the Ivorian Office of Parks and Reserves (OIPR) (the body responsible for managing two protected areas] and our research team. Data collection in Taï National Park (PNTAI) was carried out using ground surveys [15], due to the dense forest cover which limits aerial visibility. This method has the advantage of allowing for detailed detection of signs of presence, greater accuracy in species identification and the collection of detailed ecological data. Conversely, in Comoé National Park (PNCOMOE), where forest cover is sparse to open, data were obtained through aerial surveys [16]. This approach allows for the rapid coverage of large areas and provides an overall view of the distribution of animal populations. However, it may underestimate certain elusive or small-sized species and remains dependent on visibility conditions and climatic constraints. The surveys were organized as annual time series of species abundance. For each park, a ‘species’ matrix was constructed for each year, where each cell corresponds to the number of individuals recorded for a given species each year. This data structure is consistent with ecological time-series approaches used in community stability analyses [17,18].

### Analysis of community temporal dynamics

#### Total abundance

Annual total abundance was calculated as the sum of the abundances of all observed species: 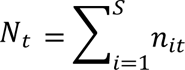 where *N_t_* represents the total abundance in year *t*, *n_it_* the abundance of species *i* in year *t*, and *S* the total number of species [17,18]. This metric allows us to assess the overall demographic changes in communities over time.

#### Alpha diversity

Species diversity was estimated using the Shannon-Wiener index, which is widely used in ecology to simultaneously account for species richness and relative abundance [19]: 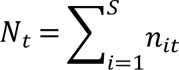 where *N_t_* represents the total annual abundance *t*, *n_it_* the abundance of the species *i* on an annual basis *t*, and *S* the total number of species. This index serves as a benchmark in studies of biodiversity and ecological stability.

#### Specific richness

Specific richness has been defined as the total number of species observed per year: S = total number of species recorded [20,21]. This measure was used to assess variations in taxonomic diversity between sites and overtime.

#### Pielou’s equity

Community equity was calculated using the Pielou index: 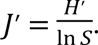 This index ranges from 0 to 1, with values close to 1 indicating a uniform distribution of abundance across species [22].

#### Assessment of functional stability

Ecological stability was analyzed across several complementary dimensions: resistance, resilience, invariance and persistence, in line with the conceptual frameworks developed by [23,24].

#### Resistance

Resistance was defined as the system’s ability to maintain its state after a disturbance: 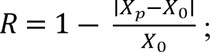 where *X*_0_ is the initial state and *X_p_* the post-disturbance state [13,23].

#### Resilience

Resilience was assessed based on the post-disturbance recovery rate: 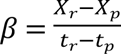 [11]. This approach allows us to quantify the rate of return to equilibrium after a perturbation.

#### Composite Stability Index

A synthetic stability index was calculated to integrate several dimensions:ICS=5R+I+P+T+[1−Syn]; where: [R] = resistance, [I] = invariance, [P] = persistence, [T] = stability, and [Syn] = synchrony [13]. This integrative approach is recommended for a multidimensional assessment of ecosystem stability.

### Analysis of dynamic stability using the Jacobian approach

The local stability of communities was studied using spectral analysis of the Jacobian matrix [25] according to the formula approach: 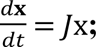 where [J] is the Jacobian matrix of interactions between species. The eigenvalues *λ_max_* were calculated to assess local stability: *Re*[*lambda*] < 0: stable system; *Re*[*lambda*] > 0: Unstable system. The dominant eigenvalue λmax was chosen as the main indicator of structural stability. This method is widely used in theoretical ecology to analyze the robustness of biological networks.

### Statistical processing and visualization

All ecological stability analyses were conducted using R [version 4.5.3; R Core Team], employing a set of complementary packages. Stability indices were calculated using the algorithms of the estar package [26], multivariate diversity analyses were performed via vegan [27], fmsb [28] to easily generate radar charts, and tabular data manipulation was carried out using dplyr [29] and tidyr [29]. The visualization relied on ggplot2 [30] for temporal dynamics and species-specific representations, patchwork [31] for assembling multi-panel Figs, and ggrepel [32] for non-overlapping annotation of key species in the recovery space. Comparisons between sites were performed on annual averages and temporal trajectories.

## Results

### Temporal dynamics of total abundance between Taï and Comoé National Parks

Analysis of the temporal dynamics of total abundance reveals contrasting trends between the two sites studied, Comoé National Park (PNCOMOE) and Taï National Park (PNTAI), highlighting marked differences in demographic regimes, ecological stability and the intensity of interannual fluctuations (Fig 2). It should be noted, however, that different data collection methods were used at these two sites: inventories at PNTAI were conducted on foot transects, whereas those at PNCOMOE were carried out via aerial surveys. This methodological difference is likely to introduce differential detectability biases between the two sites and must be taken into account in the comparative interpretation of the results.

**Fig 2.**
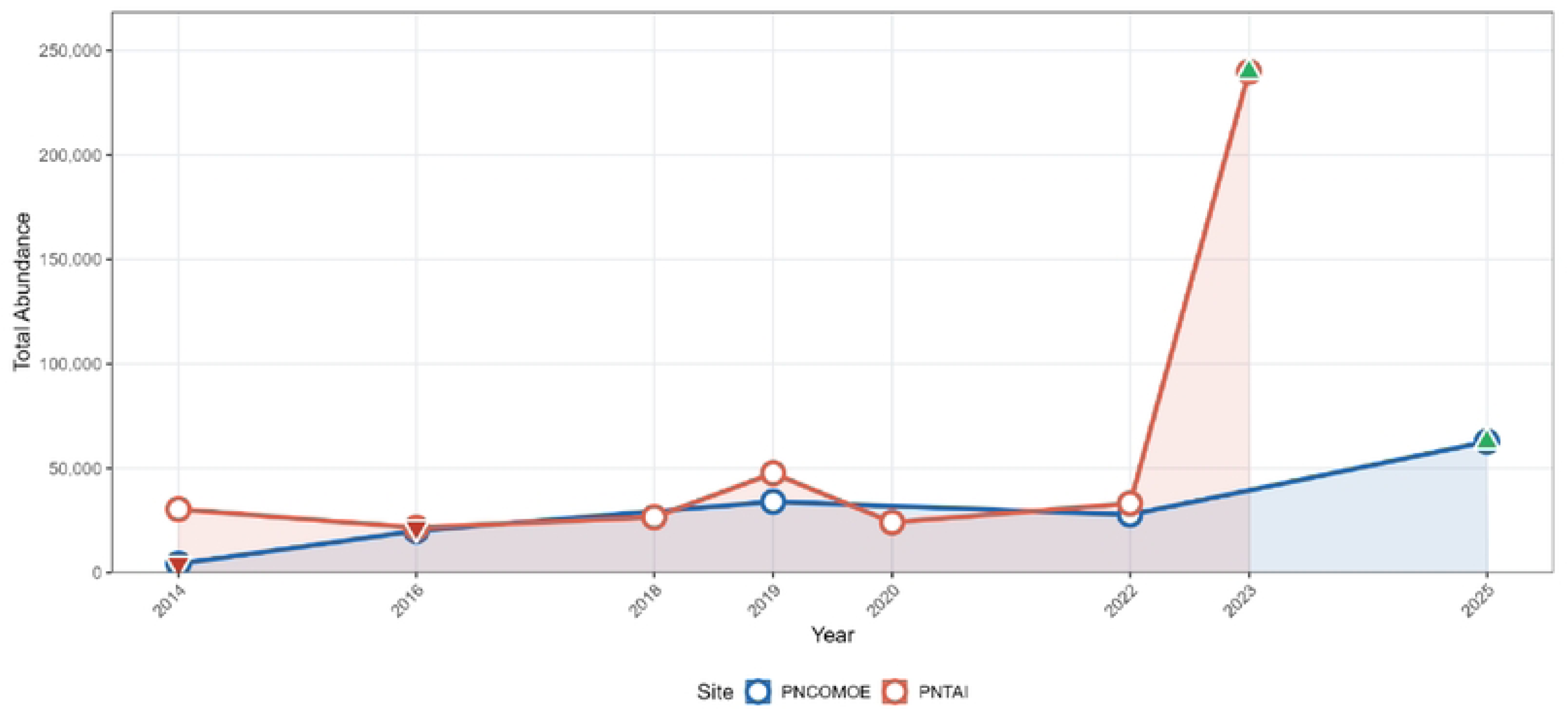
Temporal trends in total abundance in the Comoé National Park (PNCOMOE) (blue line) and Taï National Park (PNTAI) (red line) over the period 2014-2025.

At PNCOMOE, the observed dynamics reflect a gradual and relatively steady increase in total abundance over the entire period under consideration. Following an initial phase characterized by low abundance in 2014, population numbers increased significantly from 2016 onwards, then remained at relatively stable intermediate levels between 2018 and 2022, before peaking in 2025 (Fig 2). This upward trajectory suggests a positive long-term demographic trend, potentially associated with improved ecological conditions, increased availability of food resources, or enhanced effectiveness of conservation measures (Fig 2). Furthermore, the aerial surveys used in the PNCOMOE provide broader and more homogeneous spatial coverage of open savannah habitats, which helps to explain the relative consistency of abundance estimates over time. The relatively linear increase observed in the PNCOMOE also reflects strong ecological resilience, with moderate interannual variations indicating functional stability of the ecosystem.

Conversely, the PNTAI exhibits a much more unstable and highly fluctuating trend. Although initial abundances were generally higher than those of the PNCOMOE between 2014 and 2022, the population trajectory is marked by alternating significant increases and decreases, reflecting high temporal variability (Fig 2). An initial peak was observed in 2019, followed by a notable decline in 2020, then a moderate recovery in 2022 (Fig 2). However, the most striking feature remains the exceptional population explosion recorded in 2023, when the total abundance reached a level far exceeding all values observed across the entire time series (Fig 2). This abrupt peak, several times greater than previously recorded abundances, indicates either an episode of massive proliferation or a significant change in environmental conditions, such as high primary productivity, spatial clustering of individuals or massive seasonal migration. It is also possible that an intensification or standardization of the foot-trapping sampling effort contributed to this apparent increase, given that foot-trapping transects in dense forest environments are particularly sensitive to variations in effort and spatial coverage (Fig 2).

### Spatio-temporal variation in alpha diversity between Comoé and Taï National Parks

Taï National Park exhibits significantly higher diversity (H′ = 2.82 ± 0.289) than that observed in Comoé National Park (H′ = 1.57 ± 0.191), indicating a richer, more balanced and ecologically more complex biological community (Fig 3). This difference reflects distinct ecological dynamics between the two protected areas, although it is also likely to be influenced in part by the inventory protocols employed. Indeed, the foot transects used in the Taï National Park allow for more detailed detection of elusive and forest-dwelling species, particularly small mammals, primates and cryptic species, compared to the aerial surveys conducted in the Comoé National Park, which tend to favour the detection of large mammals in open habitats.

**Fig 3.**
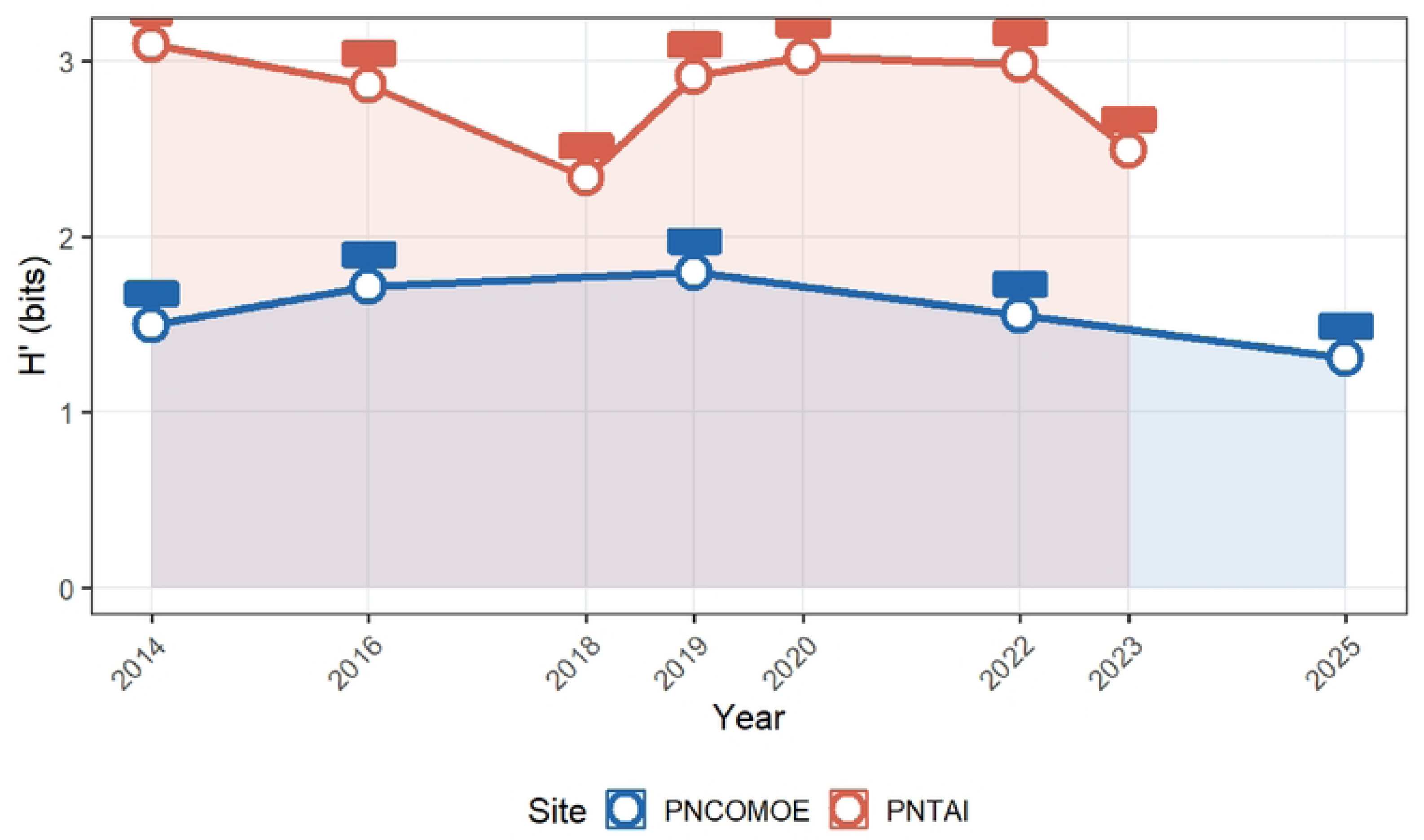
Temporal trends in alpha diversity (Shannon H′ index) between PNCOMOE and PNTAI (2014–2025).

This discrepancy is corroborated by the average species richness (S), which is significantly higher in Taï National Park (55.7 taxa) than in Comoé National Park (18.6 taxa) (Fig 3). This difference could, however, be influenced by data collection methods, as inventories were conducted via ground surveys in PNTAI, whereas aerial surveys were used in PNCOMOE. Such a difference indicates that PNTAI constitutes an ecosystem harbouring greater biological heterogeneity, likely linked to a more diverse mosaic of habitats, greater stability of ecological niches, and more complex biotic interactions favouring the coexistence of a large number of taxa. The ground-based method, which is better suited to the complex forest environment of the PNTAI, is also likely to have recorded a greater number of species with low density or reclusive behaviour, thereby contributing to higher estimates of species richness compared to aerial surveys.

The temporal analysis reinforces this ecological contrast. At PNTAI, the Shannon index remains consistently high throughout the study period, fluctuating generally between 2.3 and 3.1 bits, with a slight dip in 2018 followed by a rapid recovery between 2019 and 2022, during which time diversity reached its peak values (Fig 3). This stability at a high level reflects strong ecological resilience, the sustainable maintenance of taxonomic richness, and the robustness of biological interaction networks in the face of environmental fluctuations.

Conversely, the PNCOMOE exhibits structurally lower and more unstable alpha diversity, varying between 1.3 and 1.8 bits (Fig 3). Although a slight increase is observed between 2014 and 2019, suggesting a temporary improvement in community diversity, a gradual decline then occurs until 2025, indicating a gradual simplification of the ecological structure (Fig 3). This trend may reflect biotic homogenization, a reduction in available niches, or an increase in the competitive dominance of certain taxa, but also the inherent limitations of aerial surveys in detecting species diversity within complex savannah habitats.

### Spatio-temporal variation in species richness between PNCOMOE and PNTAI (2014-2025)

Over the entire study period, PNTAI exhibits an average species richness of 55.7 taxa, nearly three times higher than that observed at PNCOMOE (18.6 taxa) (Fig 4). This substantial quantitative difference reflects PNTAI’s significantly greater capacity to support high taxonomic diversity. However, it should be noted that this difference is partly due to the inventory methods used: the foot transects conducted at the PNTAI allow for the detection of a much broader taxonomic spectrum, including elusive forest species and small mammals, whereas the aerial surveys carried out at the PNCOMOE primarily target large mammals detectable by sight in open habitats. This methodological asymmetry must therefore be considered as a partial explanatory factor for the observed difference in species richness, in addition to the actual ecological differences between the two ecosystems.

**Fig 4.**
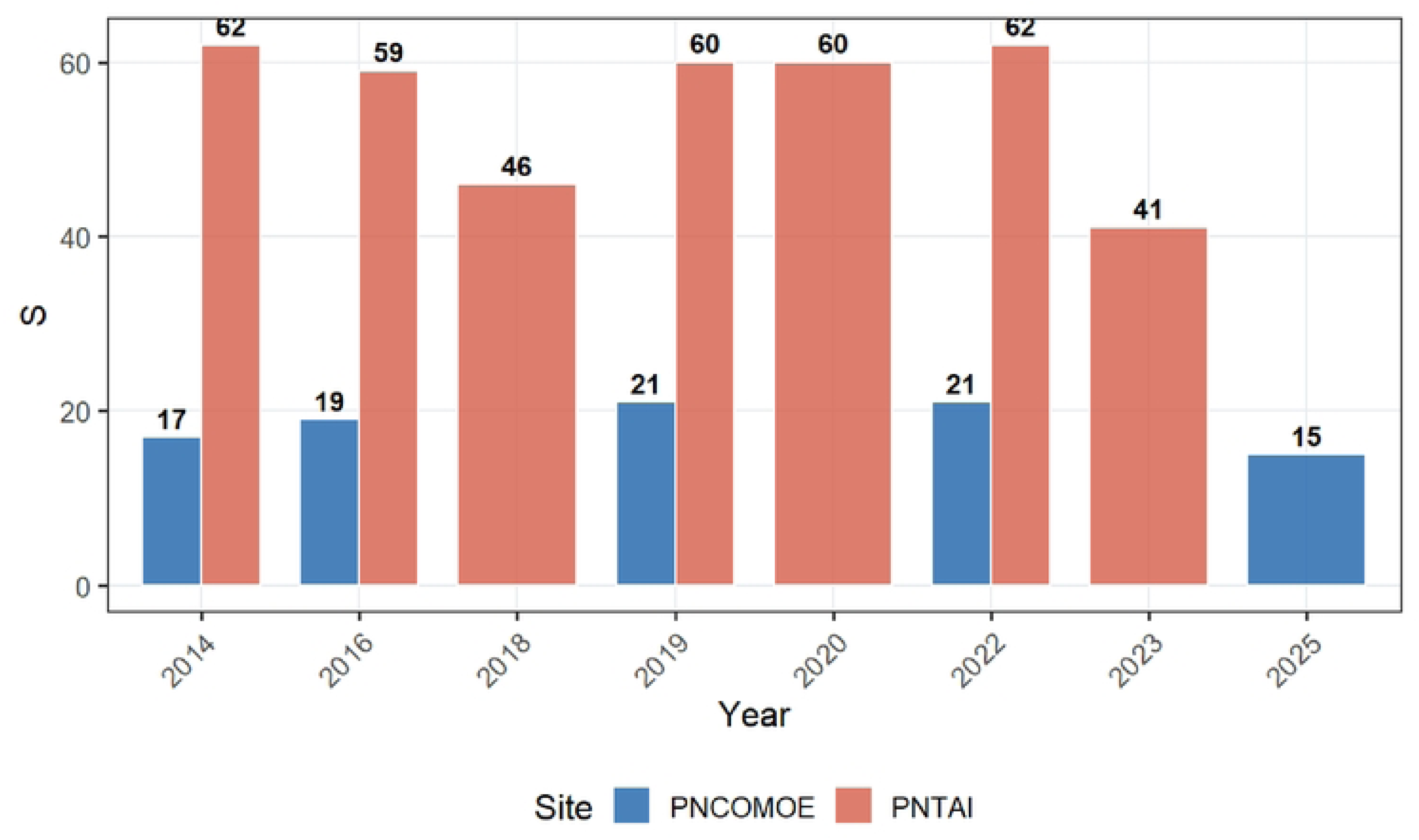
Variation temporelle de la richesse spécifique (S) entre PNCOMOE et PNTAI (2014 -2025).

Within this site, species richness remains consistently high throughout the monitoring period, fluctuating between 41 and 62 taxa (Fig 4). The peaks observed in 2014 and 2022 (62 taxa) and the sustained levels recorded in 2019 and 2020 (60 taxa) demonstrate a remarkable maintenance of taxonomic diversity, despite interannual fluctuations. Although a slight decline was observed in 2018 (46 taxa) and again in 2023 (41 taxa), levels remain well above those of the PNCOMOE, highlighting the high structural resilience of the site’s biological communities (Fig 4).

Conversely, the PNCOMOE exhibits structurally low and relatively stable species richness, ranging from 15 to 21 taxa, with an average of 18.6 taxa (Fig 4). A slight increase is observed between 2014 (17 taxa) and 2019–2022 (21 taxa), suggesting a temporary improvement in conditions favourable to taxonomic coexistence (Fig 4). However, this increase remains modest and is followed by a decline in 2025 (15 taxa), indicating a possible contraction in species diversity or a gradual homogenization of the community. It is also possible that the low taxonomic resolution of aerial surveys underestimates the actual species richness at PNCOMOE, particularly for medium-sized species that are difficult to identify from the air.

From an ecological perspective, the contrast in species richness between the two sites likely reflects major differences in habitat structure, resource availability and the complexity of biotic interactions. The high species richness at PNTAI suggests an ecosystem offering a wide variety of microhabitats and ecological niches, favouring the coexistence of a large number of taxa. Conversely, the relatively lower species richness in the PNCOMOE could reflect greater environmental constraints, higher ecological selectivity, increased dominance of certain taxa, but also the inherent detection limitations of aerial surveys.

### Temporal Dynamics of Pielou’s Evenness of Communities in the PNCOMOE and PNTAI

The Pielou Evenness index (J’) reveals different community structures between the two protected areas. The PNTAI exhibits a significantly higher average evenness (J’ = 0.701) than the PNCOMOE (J’ = 0.539), indicating that relative abundances are distributed much more homogeneously among taxa (Fig 5). This high evenness reflects low ecological dominance: no single taxon monopolizes available resources, which promotes balanced coexistence and increased functional redundancy within the community. Conversely, the relatively lower value recorded in the PNCOMOE reflects a marked abundance asymmetry, likely structured by the dominance of a limited number of opportunistic or generalist taxa, mechanically reducing the effective diversity of the community. It should be noted that aerial surveys, by preferentially detecting large, gregarious species (such as *Kobus kob* or *Alcelaphus buselaphus*), can artificially accentuate this relative abundance asymmetry at the PNCOMOE and thus lead to an underestimation of true evenness.

**Fig 5.**
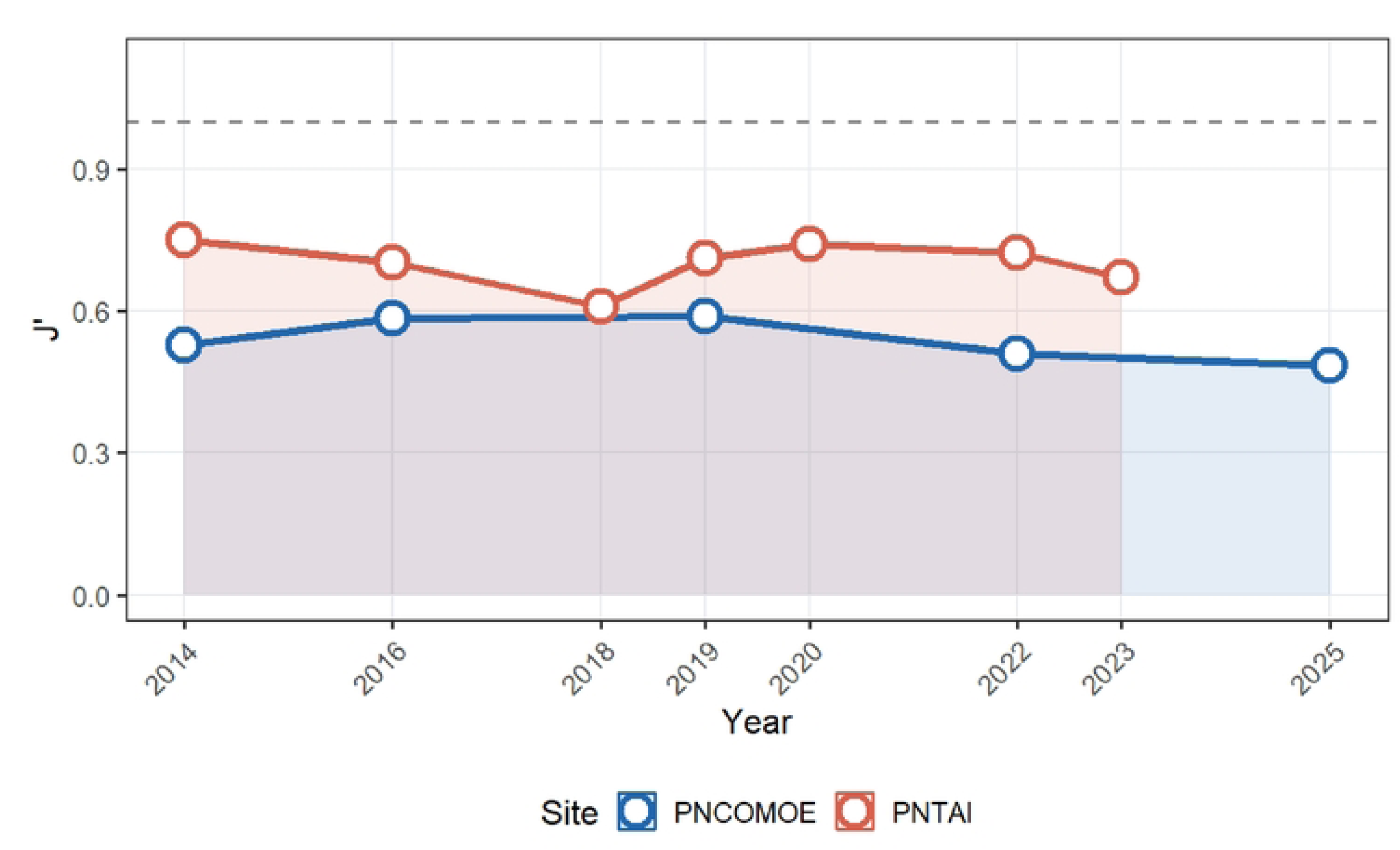
Temporal evolution of community evenness (Pielou Index J′) between PNCOMOE and PNTAI (2014-2025).

Temporally, both sites show a trend of declining evenness over the period 2014–2025, more pronounced at the PNTAI (ΔJ’ ≈ -0.10) than at the PNCOMOE (ΔJ’ ≈ -0.03) (Fig 5). This gradual convergence towards PNCOMOE values could signal ongoing community restructuring, possibly linked to environmental disturbances, differential anthropogenic pressures, or altered interspecific competition dynamics. The perfect equidistribution reference line (J’ = 1) further underlines that the two communities remain far from an ideal distribution state, which is expected in complex natural ecosystems, but reinforces the importance of the longitudinal monitoring undertaken (Fig 5).

### Analysis of functional stability metrics

These metrics were calculated from data whose structure differs according to the site and the inventory methods (pedestrian inventories in PNTAI, aerial inventories in PNCOMOE).

First, persistence reached a normalized maximum value in both cases (Persistence = 1.000), reflecting complete continuity of presence for the monitored species throughout the study period (Fig 6). This absolute structural stability indicates the absence of detectable local extinction and confirms the short-term ecological viability of the communities in both protected areas.

**Fig 6.**
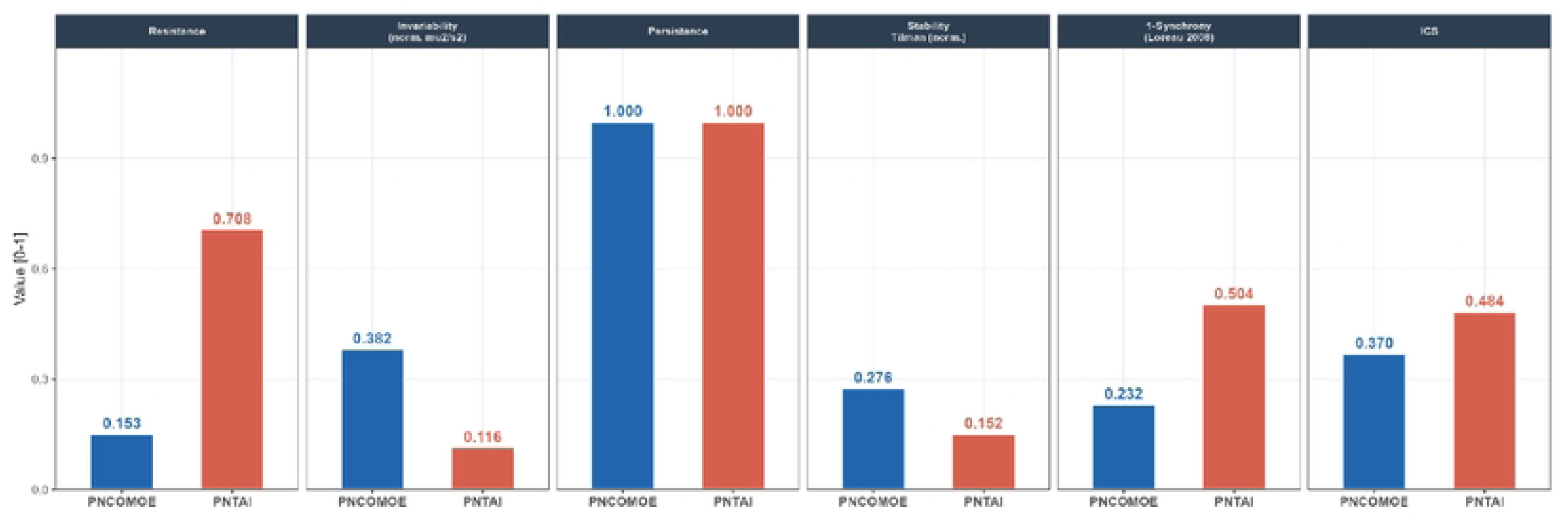
Comparison of functional stability metrics for wildlife communities between Comoé and Taï National Parks.

However, the communities’ capacity to absorb immediate disturbances differs significantly between the two sites. PNTAI exhibits exceptionally high resilience (0.708), 4.6 times greater than that observed at PNCOMOE (0.153) (Fig 6). This result reveals a strong functional robustness of the biological assemblages at PNTAI, capable of maintaining their abundances despite environmental fluctuations or external pressures. Conversely, the low resilience recorded at PNCOMOE suggests increased sensitivity of the population to disturbances, reflecting a more pronounced ecological vulnerability to climatic or anthropogenic shocks. Nevertheless, aerial surveys, by capturing only a fraction of the variations in the abundance of forest or small-sized species, may lead to an underestimation of the true resilience of the PNCOMOE.

Paradoxically, this lower resilience of the PNCOMOE is accompanied by a significantly higher invariability (0.382 versus 0.116 in the PNTAI) (Fig 6), indicating a more regular and predictable temporal dynamic of aggregate abundances. In other words, although more sensitive to localized disturbances, the PNCOMOE community maintains a more stable ecological trajectory over the long term, characterized by small interannual fluctuations. This trend is confirmed by the normalized Tilman stability, which is also higher in the PNCOMOE (0.276 versus 0.152 in the PNTAI) (Fig 6), reflecting greater consistency of community biomass over time. The regularity of aerial surveys and the nature of the targeted species, primarily large herbivores with slow population dynamics, may also contribute to the relative apparent stability of abundance estimates in the PNCOMOE.

Conversely, the PNTAI appears to leverage another major source of stability: interspecific asynchrony. The Loreau (2008) 1-synchrony index reaches 0.504 there, compared to only 0.232 in the PNCOMOE (Fig 6), revealing more asynchronous abundance fluctuations between species. This high degree of asynchrony constitutes a powerful ecological compensation mechanism; when some species decline, others increase simultaneously, thus dampening the overall variability of the system. This "ecological portfolio" phenomenon is widely recognized as one of the main drivers of functional stability in rich and complex ecosystems.

### Analysis of stability trajectories between Comoé and Taï National Parks

The radar chart of the five dimensions of functional stability (resistance, normalized Tilman stability, recovery, composite stability index (CSI), and Loreau’s 1-synchrony, 2008) highlights two clearly different ecological stability profiles between Comoé and Taï National Parks (Fig 7). Neither park dominates all indicators, demonstrating that each possesses its own mechanisms of ecological stability and resilience. However, caution is warranted in the comparative interpretation, as the observed differences result from a combination of actual ecological factors and potential biases related to the distinct inventory methods used at each site: foot transects for Taï National Park and aerial surveys for Comoé and Taï National Parks.

**Fig 7.**
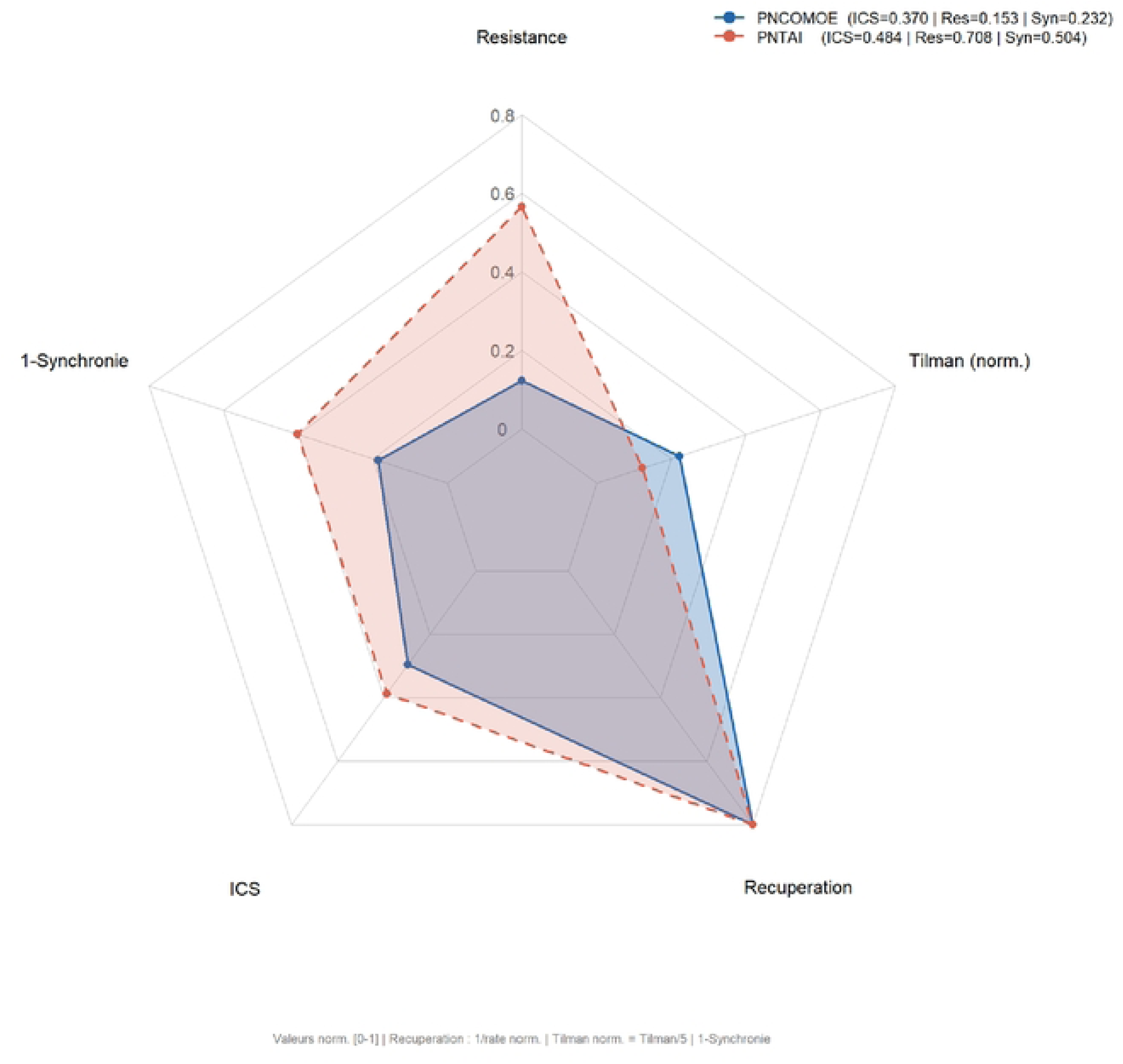
Ecological stability profile of the faunal communities of Comoé and Taï National Parks, estimated using the radar chart of indices.

First, the resistance axis represents the most marked divergence between the two parks. PNTAI exhibits a significantly higher value (0.708), forming the outermost extension of the radar polygon, while PNCOMOE remains strongly contracted towards the center (0.153) (Fig 7). This difference reflects a significantly greater capacity of PNTAI communities to maintain their abundance structure in the face of immediate disturbances, confirming high functional robustness.

Second, the 1-synchrony axis, representing the asynchrony of interspecific fluctuations, constitutes a second major gradient of differentiation. PNTAI stands out with a higher value (0.504 versus 0.232 in PNCOMOE), indicating a greater desynchronization of population dynamics between species (Fig 7). This pattern favors a functional compensation effect, a mechanism by which the temporary decline of some species is offset by an increase in others, thus stabilizing the overall biomass of the ecosystem.

Conversely, the PNCOMOE exhibits a relative advantage on the normalized Tilman stability axis (0.276 versus 0.152 for the PNTAI), suggesting lower temporal variability in aggregate biomass and more consistent long-term community dynamics (Fig 7). This signature reflects stability based more on the consistency of ecological flows than on the capacity to absorb disturbances. The regularity of the aerial survey protocols also contributes to reducing the estimated variance of aggregate biomass.

Furthermore, both polygons converge towards their maximum on the recovery axis (1.000), the only axis of perfect symmetry in the diagram (Fig 7). This convergence indicates complete persistence of the monitored species, with no detectable local extinctions, attesting to remarkable ecological continuity in both protected areas.

Finally, the Composite Stability Index (CSI) confers a moderate overall advantage to the PNTAI (0.484 versus 0.370 for the PNCOMOE) (Fig 7). However, the polygon’s geometry clearly shows that this superiority is not uniform across all dimensions, but is primarily based on the synergistic combination of its strong resilience and significant interspecific asynchrony.

### Dynamic stability by Jacobian analysis of communities

Local stability analysis using the Jacobian method reveals that the two studied ecosystems exhibit an unstable dynamic regime over the period 2016-2025. The dominant eigenvalues of the community matrix are positive for both sites: Re(λ_max) = 3.73 for Comoé National Park and Re(λ_max) = 5.58 for Taï National Park, indicating that disturbances applied at equilibrium are not dampened but amplified over time (Fig 8). Although many of the eigenvalues have negative real parts, reflecting partial self-regulation of interspecific interactions, the presence of a strictly positive dominant eigenvalue is sufficient to invalidate the overall stability of the system.

**Fig 8.**
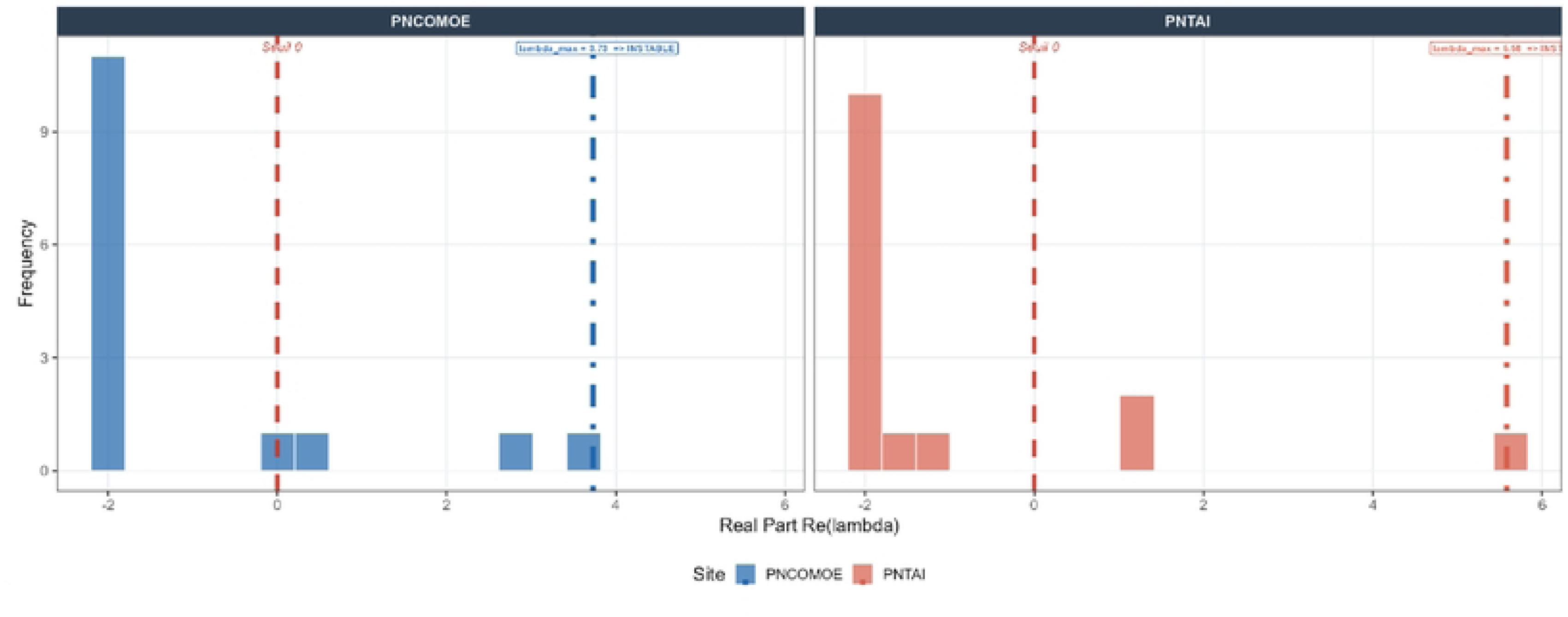
Local stability of wildlife communities assessed by spectral analysis of the Jacobian matrix in Comoé and Taï National Parks (2014-2025).

The inter-site comparison reveals a significant instability gradient: the PNTAI (5.58) exhibits a dominant eigenvalue 49.6 % higher than that of the PNCOMOE (3.73), reflecting a more pronounced structural vulnerability of the Taï Park community (Fig 8). This difference could reflect higher trophic connectance, greater taxonomic heterogeneity, or more intense habitat pressures within the PNTAI. However, it is important to note that the Jacobian matrix is constructed from abundance time series whose resolution depends on the inventory protocol. The PNTAI foot transects capture richer taxonomic diversity and temporal fluctuations, which can amplify the interactions estimated in the matrix and, consequently, increase the dominant eigenvalue. The aerial inventories of the PNCOMOE, by smoothing out some of the variability of species with low aerial detectability, tend to produce more condensed and potentially less unstable- looking community matrices.

These results have direct implications for conservation: an unstable Jacobian ecosystem is theoretically susceptible to abrupt regime shifts in response to even small-scale disturbances. Management strategies aimed at strengthening stabilizing interactions, particularly mutualistic effects and top-down regulation, therefore appear to be priorities for both sites, with increased urgency for the PNTAI.

### Resistance and persistence of dominant wildlife species in two protected areas (2014-2025)

The combined analysis of resistance and persistence reveals a highly asymmetrical distribution of stability capacities within the wildlife communities of the two protected areas. In both sites, most species exhibit a resistance below 0.10 (Fig 9), reflecting widespread vulnerability to environmental and anthropogenic disturbances. Only a small core of species stands out with high resistance values, thus defining the key species for community stability dynamics.

**Fig 9.**
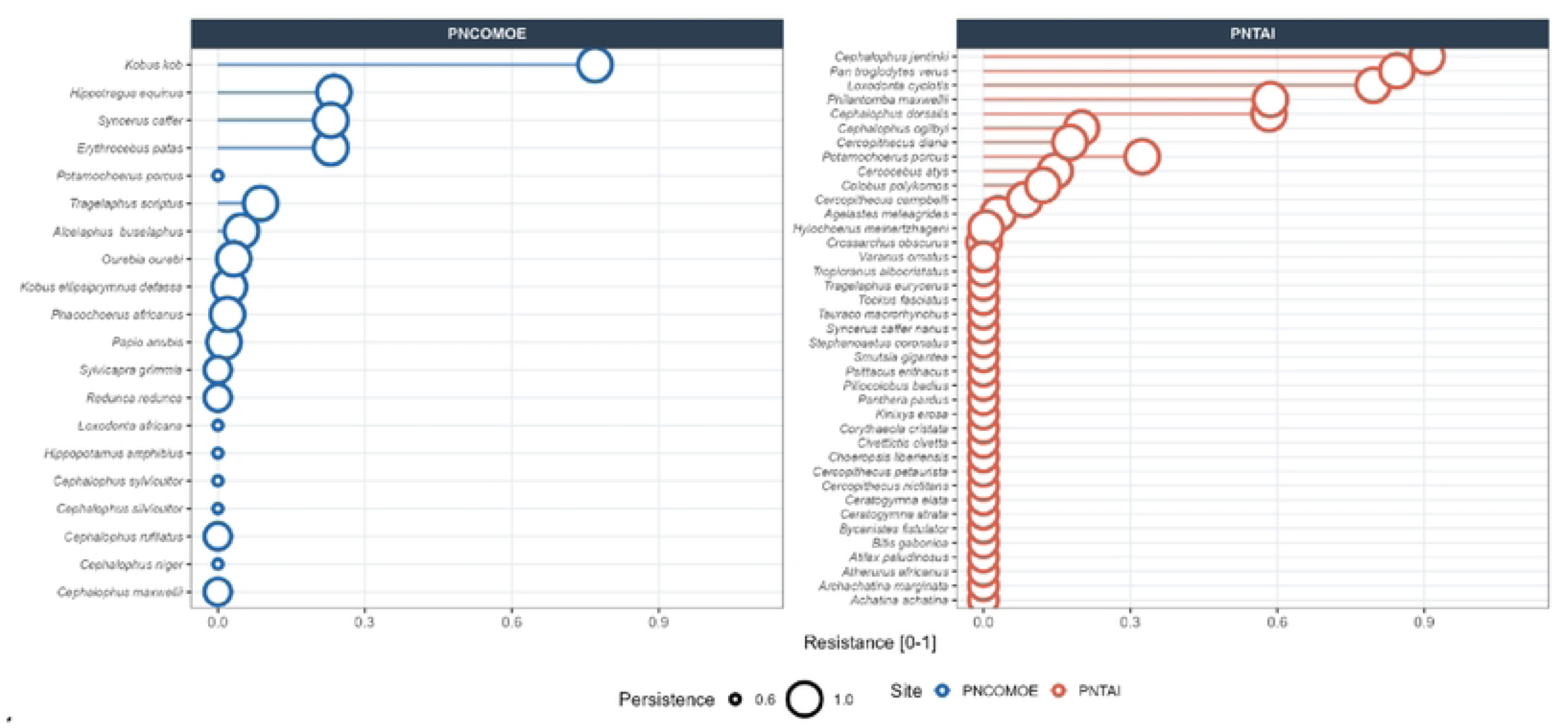
Distribution of resistance and persistence among the 20 most contributing species within the wildlife communities of Comoé and Taï National Parks.

In Comoé National Park, Kobus kob clearly stands out as the species with the highest resistance (Re ≈ 0.85), followed at a distance by Hippotragus equinus and Syncerus caffer (Fig 9). This profile, dominated by large savanna herbivores, reflects the trophic structure characteristic of the Comoé River’s Sudanian-Guinean ecosystems, where these species play a key role in maintaining energy flows and regulating vegetation. It is worth noting that the aerial surveys used at the Comoé National Park are particularly effective at estimating the abundance of these large herbivores in open areas, which is reflected in their estimated high resilience. Conversely, the resilience of smaller or less conspicuous forest species, which are potentially less easily detected by aerial surveys, may be underestimated in this analysis.

In the PNTAI, the resistance profile is more diverse and higher: *Cephalophus jentinki* (Re ≈ 0.95), *Pan troglodytes verus* (Re ≈ 0.90), and *Loxodonta cyclotis* (Re ≈ 0.85) are the three highly resistant species, in addition to several duiker species with intermediate resistance (0.3–0.6) (Fig 9). This abundance of resistant species reflects the structural complexity of the dense humid forest of Taï and the functional diversity of the guilds that compose it. The presence of the chimpanzee (*Pan troglodytes verus*) and the forest elephant (*Loxodonta cyclotis*), two ecosystem engineers, among the most resistant entities underscores their irreplaceable role in the resilience of this ecosystem. Pedestrian transects allow for the precise detection of these species, even in a dense forest environment, thus strengthening the robustness of resistance estimates in the PNTAI.

From a conservation perspective, these results argue for species-centered and differentiated management strategies: in the PNCOMOE, the priority protection of *Kobus kob* and large savanna ungulates constitute the main lever for maintaining community stability. In the PNTAI, the simultaneous conservation of *Pan troglodytes verus*, *Loxodonta cyclotis*, and rare forest duikers (*C. jentinki, C. dorsalis*) appear essential to preserve the functional architecture of this ecosystem, a UNESCO World Heritage Site (Fig 9).

### Ecological Resilience and Recovery Capacity of Mammalian Communities in Comoé and Taï National Parks

Analysis of the post-disturbance recovery space reveals radically divergent resilience trajectories between the two protected areas, both at the community and interspecific levels. In Comoé and Taï National Parks, community resilience is undefined (RE = NA), which constitutes a critical ecological signal: in the absence of a sufficiently dominant negative eigenvalue, the system lacks a formally calculable return-to-equilibrium trajectory. Nearly all species exhibit a zero or negative β recovery rate, and only *Sylvicapra grimmia* (RE ≈ 2.0, low β) and *Cephalophus rufilatus* (RE ≈ 1.1) reach or exceed the threshold for complete recovery (Fig 10). The projected duration of 12,467 days (∼34 years) reflects not actual recovery but the residual persistence of a system in decline (Fig 10). These results indicate an ecosystem under chronic stress, likely subjected to cumulative pressures (habitat degradation, poaching, fragmentation) that have eroded its intrinsic resilience. Furthermore, aerial surveys, by failing to detect certain small or forest species with high potential resilience, may lead to an underestimation of the actual overall resilience of the PNCOMOE.

**Fig 10.**
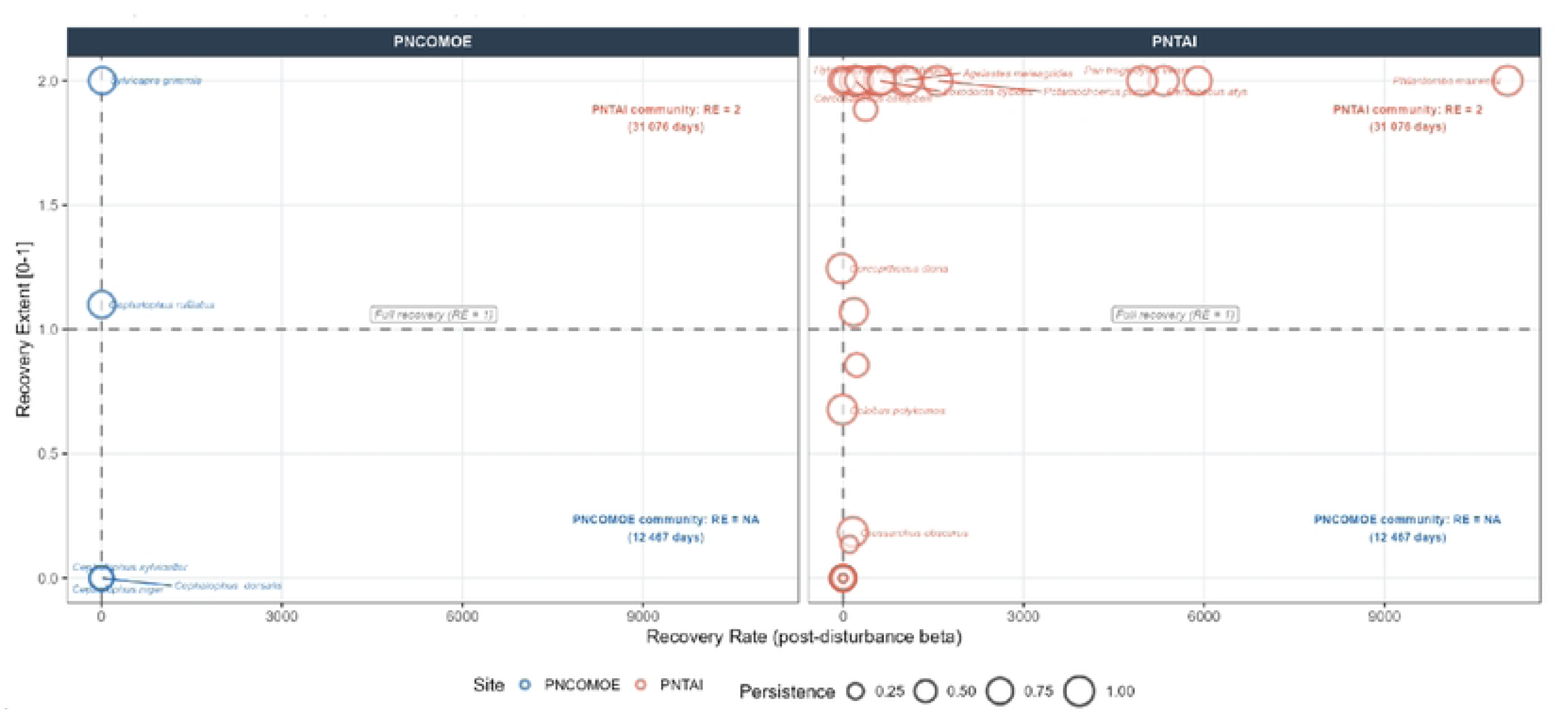
Post-disturbance recovery area of wildlife communities in Comoé and Taï National Parks (2014-2025).

At the PNTAI, the picture is structurally more favorable but not without concerns. Community resilience is defined and high (RE = 2), meaning that the community can theoretically recover beyond its pre-disturbance state, a phenomenon that can be interpreted as ecological overcompensation characteristic of complex, highly connected forest systems (Fig 10). A dense core of species including *Loxodonta cyclotis, Pan troglodytes verus*, *Philantomba maxwellii, Potamochoerus porcus*, and several duikers simultaneously exhibits high β recovery rates (up to ∼9,000) and recovery extents greater than 1, with moderate to high persistence (Fig 10). However, the projected community duration of 31,076 days (∼85 years) reveals the temporal cost of this resilience: even in the most favorable scenario, the return to equilibrium occurs on a transgenerational timescale, incompatible with conventional protected area management cycles (Fig 10).

Several species in the PNTAI occupy intermediate positions: *Cercocebus atys, Colobus polykomos*, and *Crossarchus obscurus* show recovery areas between 0 and 1 with low β values, indicating partial and slow recovery (Fig 10). *Crossarchus obscurus*, positioned at RE ≈ 0 and β ≈ 0, represents a borderline case analogous to the most vulnerable species in the PNCOMOE (Fig 10).

### Temporal dynamics of wildlife abundance, stability of dominant species, erosion of sensitive species, and early signs of community collapse in Comoé and Taï National Parks

Temporal analysis of wildlife abundance using heatmaps reveals contrasting community dynamics between the two protected areas over the decade 2014-2025, with concerning warning signs in both systems (Fig 11).

**Fig 11.**
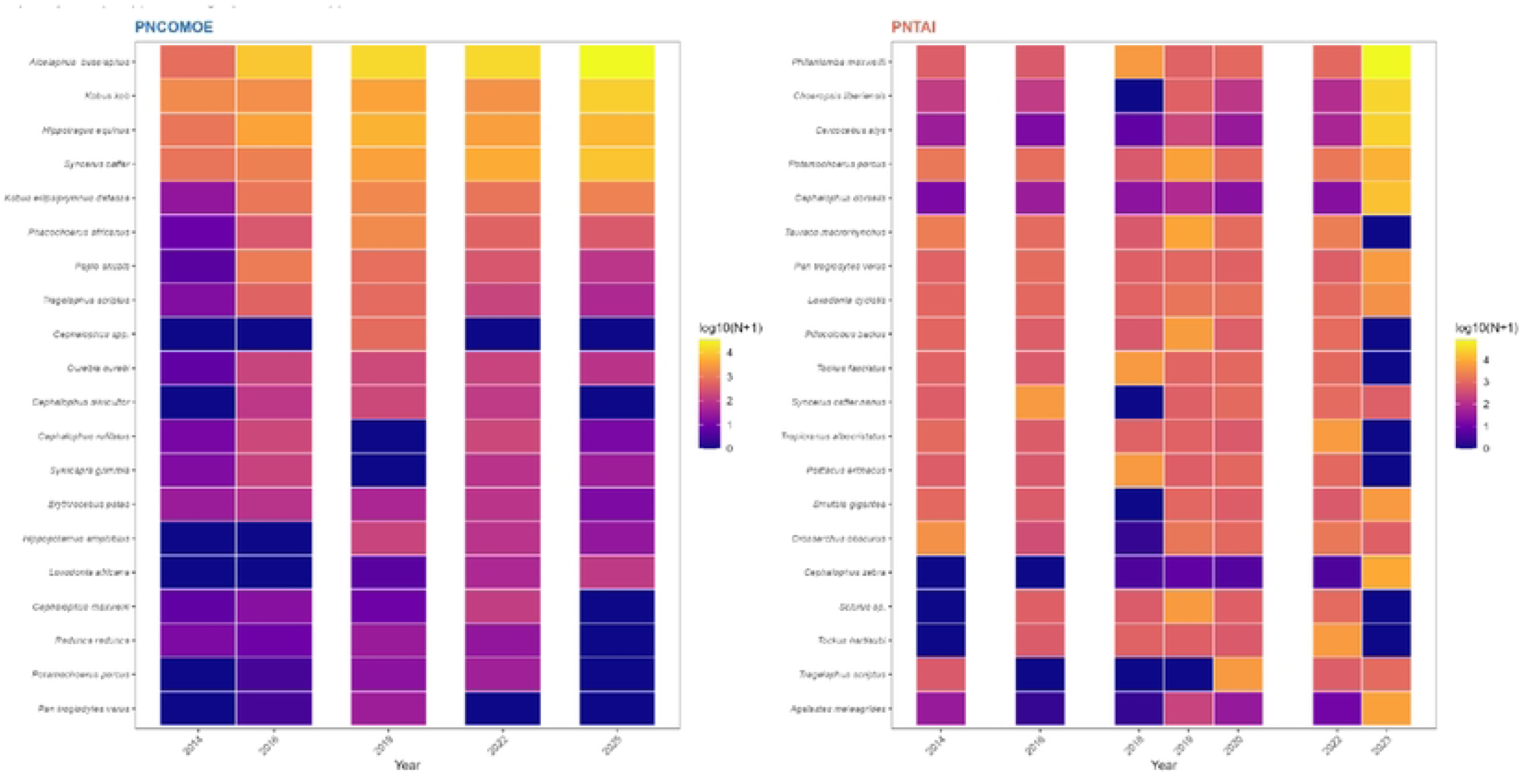
Temporal dynamics of the abundance of the 20 dominant wildlife species in two protected areas of Côte d’Ivoire (2014-2025).

In the PNCOMOE, the large savanna herbivore species *Alcelaphus buselaphus, Kobus kob, Hippotragus equinus*, and *Syncerus caffer* maintained high and relatively stable abundances over the entire period [log₁₀(N+1) ≥ 3–4], confirming their status as key species in this Sudanian-Guinean ecosystem (Fig 11). These species are precisely the optimal targets for aerial surveys in open savanna environments, which strengthens the reliability of the estimates for these taxa. This stability of the dominant species contrasts, however, with a progressive decline in low-abundance species: *Cephalophus rufilatus, Sylvicapra grimmia, Hippopotamus amphibius,* and *Loxodonta africana* exhibit increasingly cooler color signals over time, reflecting a continuous decline (Fig 11). At PNTAI, the temporal dynamics are marked by more pronounced spatio-temporal heterogeneity, reflecting the functional complexity of the forest ecosystem. The dominant species *Philantomba maxwellii, Choeropsis liberiensis, Cercocebus atys*, and *Potamochoerus porcus* maintain high abundance levels over most of the period, although significant interannual fluctuations are noticeable (Fig 11). The 2019–2022 period marks a dynamic turning point for several species: *Loxodonta cyclotis, Tauraco macrorhynchus*, and *Pan troglodytes verus* show notable declines, before an apparent stabilization or slight recovery in 2025 for some (Fig 11). These potential local extinctions specifically concern specialist forest species with low tolerance to disturbances, which are particularly well detected by pedestrian transects; their disappearance therefore constitutes a reliable and sensitive indicator of ecosystem integrity degradation.

In the future, integrating these abundance heatmaps into a real-time monitoring system coupled with ecological stability indices would make it possible to identify locally endangered species before their extinction is complete. Furthermore, harmonizing protocols between sites would strengthen data comparability and the robustness of ecological assessments.

## Discussion

### Divergence of population trajectories between actual ecological signature and methodological artifact

The contrasting abundance trajectories between the PNTAI and the PNCOMOE constitute the first key finding of this study. Such inter-site divergence in the temporal dynamics of wildlife populations is widely documented in the African literature on protected areas and generally reflects the combination of factors intrinsic to ecosystems [habitat structure, resource availability, disturbance regimes] and extrinsic factors related to differential anthropogenic pressures [33,34]. The gradual and steady growth observed in the PNCOMOE is typical of large savanna herbivore communities under effective protection and relatively stable climatic conditions [35]. Conversely, the pulsating and strongly non-linear dynamics of the PNTAI are reminiscent of fluctuation patterns characteristic of tropical rainforests, where primary productivity, resource phenology, and the complexity of biotic interactions generate rapid demographic responses that are often disproportionate to disturbances [36,37].

However, aerial surveys used in the PNCOMOE offer homogeneous spatial coverage and are recognized as particularly well-suited to detecting large gregarious mammals in open environments [38,39], but are limited by reduced detectability for small, solitary, forest-dwelling, or intermediate-sized species [40]. Conversely, pedestrian transects conducted in the PNTAI according to distance sampling protocols [15] allow for a more precise and comprehensive estimate of taxonomic diversity in forest environments but are sensitive to variations in sampling effort and the spatial heterogeneity of vegetation cover [41]. The exceptional peak in abundance recorded at PNTAI in 2023 precisely illustrates this ambiguity: while it may reflect a genuine ecological episode of proliferation or spatial regrouping, it could just as easily result from an intensification of fieldwork efforts or a change in the collection protocol, a bias commonly reported in time series of wildlife inventories [42,43]. These considerations underscore the critical need to harmonize inventory protocols between sites in long-term comparative studies, in accordance with the international recommendations of the TEAM (Tropical Ecology Assessment and Monitoring Network) and the IUCN standards for biodiversity monitoring [44].

### Alpha Diversity Differentiation

The considerable difference in alpha diversity between the PNTAI (H′ = 2.82) and the PNCOMOE (H′ = 1.57) is consistent with the well-documented diversity gradients between the dense humid forests and the Guinean savannas of West Africa [45,46]. The Taï Forest, recognized as one of the last great stands of primary forest in West Africa and a UNESCO World Heritage Site, constitutes a global biodiversity hotspot whose exceptional species richness is widely established [47,48]. The lower diversity of the PNCOMOE reflects the structural characteristics of the Sudanian-Guinean ecosystems, dominated by transitional savanna-forest-gallery vegetation, offering a more limited range of ecological niches for terrestrial megafauna [49].

Nevertheless, the observed difference in species richness 55.7 taxa in the PNTAI versus 18.6 in the PNCOMOE exceeds what ecological differences between biomes alone can fully explain and calls for cautious interpretation. Foot surveys using distance sampling allow for the detection of elusive, nocturnal, or low-biomass species, particularly small duikers, arboreal primates, and rodents, which are systematically under-detected during aerial surveys [40,50]. In this context, the higher biodiversity of the PNTAI reflects both an ecological reality the structural complexity of the tropical rainforest and a methodological advantage linked to the greater sensitivity of pedestrian transects for capturing the diversity of cryptic species. This analytical distinction is fundamental to avoiding overestimating inter-site ecological differences in protected area management interpretations [43,51].

The progressive erosion of Pielou’s evenness at both sites over the period 2014-2025 (ΔJ′ ≈ -0.10 at the PNTAI; ΔJ′ ≈ -0.03 at the PNCOMOE) warrants particular attention. This signal, consistent with global trends in biotic homogenization documented in African protected areas under increasing pressure [52,53], reflects a restructuring of communities favoring dominant generalist species at the expense of sensitive specialist species. This process of eroding evenness, when it intensifies over time, constitutes a reliable early indicator of ecological degradation [13,54].

### Functional stability of ecosystems [comparison of resistance and interspecific compensation mechanisms]

The PNTAI exhibits dynamic stability based on strong resistance to disturbances (0.708) and high interspecific asynchrony (1-synchrony = 0.504), while the PNCOMOE displays conservative stability based on temporal regularity (invariance = 0.382) and the constancy of aggregate biomass (Tilman stability = 0.276).

This dichotomy fits perfectly within the theoretical framework developed by [55], according to which the stability of ecological communities can be broken down into two fundamental components: community-level resilience, which depends on the average resilience of individual populations, and interspecies compensation, which results from the asynchrony of their temporal fluctuations. The strong asynchrony observed at PNTAI has an "ecological portfolio effect" [56]. They show that this buffering mechanism allows rich and complex communities to dampen global variability despite significant individual fluctuations. This result is consistent with the theory predicting that biological diversity, by increasing the probability of asynchrony between species, constitutes one of the main guarantors of the functional stability of ecosystems [57].

Conversely, the more conservative stability of the PNCOMOE, based on the regularity of aggregate biomass rather than on active compensation mechanisms, is characteristic of low-diversity communities dominated by a few abundant species [58]. In these systems, overall stability depends more on the dynamics of dominant species, here *Kobus kob, Alcelaphus buselaphus*, and *Syncerus caffer*, than on interspecific compensation effects, making them structurally more vulnerable to the disappearance or decline of keystone species [59]. The low resistance to disturbances recorded in the PNCOMOE (0.153) confirms this vulnerability and suggests that moderate environmental or anthropogenic shocks could lead to disproportionate abundance disturbances in this ecosystem.

The convergence towards maximum persistence (1,000) at both sites nevertheless constitutes a positive ecological signal, indicating the absence of detectable local extinction over the study period. This result, while encouraging, must be interpreted with caution because the persistence measured here reflects the presence of the monitored species, and not necessarily the maintenance of their long-term viability [60]. Populations maintained at very low numbers, as appears to be the case for *Pan troglodytes verus* at PNCOMOE, can exhibit apparent persistence while being functionally extinct in terms of their ecological role [59].

### Jacobian Instability as a Systemic Warning Sign

Local stability analysis using the Jacobian method reveals that both ecosystems operate within a dynamically unstable regime, with dominant positive eigenvalues (Re(λmax) = 3.73 for PNCOMOE; Re(λmax] = 5.58 for PNTAI). This result, seemingly counterintuitive given the maximum persistence indicators, is actually consistent with the predictions of the stability theory of complex ecological networks [44]. [61] mathematically demonstrated that the stability of ecological communities decreases with increasing trophic connectance and the number of species, due to the multiplication of potentially destabilizing interactions. The higher dominant eigenvalue in PNTAI [49.6% higher than that of PNCOMOE] directly reflects its greater taxonomic richness and higher trophic connectance, consistent with the properties of tropical forest networks [62].

The presence of positive dominant eigenvalues in both parks indicates that these systems do not spontaneously return to equilibrium after disturbance [63]. This regime of local instability, however, is not synonymous with imminent collapse: many complex natural ecosystems operate permanently far from their theoretical equilibrium, maintained in quasi-stable states by feedback mechanisms [64]. The detection of Jacobian instability is therefore a first-order warning signal, indicating the need for enhanced monitoring and preventive measures to avoid irreversible shifts [65].

### Keystone Species, Ecosystem Engineers, and the Architecture of Community Resilience

The identification of a small core of highly resilient species at each of the two sites, *Kobus kob* at PNCOMOE; and *Cephalophus jentinki, Pan troglodytes verus*, and *Loxodonta cyclotis* at PNTAI directly relates to the concept of keystone species and ecosystem engineers, whose disproportionate role in maintaining the structure and function of ecosystems is now firmly established [67]. At PNTAI, the simultaneous presence of the forest elephant (*Loxodonta cyclotis)* and the chimpanzee (*Pan troglodytes verus)* among the most resilient species is particularly significant. These two species are recognized as major ecosystem engineers: *Loxodonta cyclotis* physically structures the forest habitat by creating gaps and dispersing seeds from large trees, contributing to canopy renewal [68]; *Pan troglodytes verus* exerts top-down regulation on fruit populations and ensures irreplaceable zoochorous dispersal for many plant species [69]. The functional loss of these engineer species would trigger a cascade of structural reorganizations throughout the food web, compromising the recovery capacity of the entire community [59].

In the PNCOMOE, the functional dominance of *Kobus kob* in the community stability architecture reflects its central role in the energy flows of the Sudanian-Guinean ecosystems. This species, which frequently forms large herds in humid savanna, is the primary prey of many large predators [cheetah, lion, leopard], and its population dynamics directly influence the structure and stability of the local food web [70]. Its high resilience (Re ≈ 0.85) suggests that the density-dependent regulatory mechanisms specific to this species, particularly resource regulation during the dry season, provide the system with a form of demographic buffer against disturbances. However, the concentration of community resilience on a very small number of species simultaneously constitutes a major structural weakness: the disappearance or severe decline of *Kobus kob* would expose the entire food web to a risk of cascading instability [36].

### Community Resilience and Critical Thresholds of Functional Degradation

The most critical divergence between the two sites lies in their post-disturbance resilience capacities: a defined and high resilience at PNTAI (RE = 2) versus an indefinite resilience at PNCOMOE (RE = NA). This contrast reflects fundamentally distinct functional states according to the theory of ecological resilience [65]. The lack of quantifiable community resilience at PNCOMOE is a particularly worrying signal, suggesting that this park operates within a low-capacity "catchment basin," characteristic of systems that have crossed or are approaching a critical threshold of functional degradation [71]. In this type of configuration, the system gradually loses its intrinsic capacity to absorb disturbances and return to its reference state, becoming increasingly vulnerable to abrupt and irreversible regime shifts even in response to low-amplitude disturbances [71].

The projected recovery time of ∼34 years at PNCOMOE does not reflect a return-to-equilibrium trajectory but rather the residual drift of a system without a stable attractor, a crucial distinction for protected area management. Conversely, the high resilience of PNTAI [RE = 2] and the recovery time of ∼85 years demonstrate a system capable of ecological overcompensation after disturbance, a characteristic of complex tropical forests with high functional connectivity [66]. However, a recovery time of 85 years falls within a transgenerational timescale, incompatible with conventional conservation policy planning horizons [5-20 years], which underscores the urgency of proactive rather than reactive management [72].

### Implications for the Conservation and Differentiated Management of Protected Areas

The results of this study argue for a differentiated conservation approach, adapted to the specific resilience mechanisms of each protected area. For the PNCOMOE, the data converge to indicate a state of advanced functional vulnerability, marked by low resistance, indefinite resilience, and signs of collapse for emblematic species. Active restoration interventions reintroduction or reinforcement of key species populations, restoration of ecological corridors linking the park to adjacent gallery forests and strengthened anti-poaching efforts appear urgent to halt the current trajectory of degradation [74]. The management of the PNCOMOE should also rely on real-time monitoring mechanisms for early warning signals, such as increased temporal variance in abundances, erosion of temporal autocorrelation, and critical slowing down of population dynamics [75].

For the PNTAI, whose functional stability depends on maintaining interspecific asynchrony and the resilience of ecosystem engineer species, the absolute priority is the conservation of *Loxodonta cyclotis* and *Pan troglodytes verus*, whose irreplaceable ecosystem functions determine the resilience of the entire community. Protecting these species involves not only reducing direct pressures (poaching, illegal capture), but also maintaining connectivity between the PNTAI and surrounding forest ecosystems, which is essential for maintaining viable metapopulations and the genetic diversity of populations [75].

Finally, from a methodological standpoint, this study strongly underscores the need to harmonize inventory protocols in long-term comparative monitoring programs for Ivorian protected areas. Adopting multi-method protocols combining pedestrian distance sampling, aerial surveys, and camera traps, such as those recommended by the TEAM network [44] or the IUCN/SSC standards, would represent a major step forward for the robustness of biodiversity assessments in West Africa.

## Conclusion

The PNTAI and the PNCOMOE illustrate two contrasting trajectories of ecological stability. The PNTAI exhibits dynamic stability, based on high resilience (0.708) and high interspecific asynchrony (Loreau index = 0.504), whereas the PNCOMOE is based on conservative stability, characterized by temporal regularity in abundance (invariability = 0.382) and biomass constancy (Tilman stability = 0.276). The ICS confirms the overall advantage of the PNTAI (0.484 versus 0.370). The interpretation of these contrasts must, however, take methodological biases into account: the ground surveys in the PNTAI favour the detection of cryptic species, whereas the aerial surveys in the PNCOMOE, which are effective for large mammals in open habitats, result in variable detection probabilities. The observed differences therefore reflect both ecological realities and methodological artefacts. Despite these nuances, signs of structural vulnerability converge: the absence of calculable resilience and positive Jacobian eigenvalues indicate locally unstable dynamics. At PNCOMOE, conservation efforts should target *Kobus kob, Hippotragus equinus* and *Syncerus caffer*; in the PNTAI, the conservation of *Loxodonta cyclotis, Pan troglodytes verus* and rare duikers remains essential. Projected recovery trajectories of approximately 85 years for the PNTAI, and indeterminate for the PNCOMOE, underscore the urgency of current decisions.

## Acknowledgments

We would like to thank the Office Ivoirien des Parcs et Réserves (OIPR) for granting access to the protected areas and for their contribution to data collection. Finally, we thank the local communities living near the studied forests and the field guides for their invaluable collaboration.

